# NeuroMetaBench: Expert-curated benchmark for evaluating automated neuroimaging meta-analysis

**DOI:** 10.64898/2026.11.20.768226

**Authors:** Alejandro de la Vega, James D. Kent, Anthony Yang, Shaliha C. Guarnera, Riya Louis, Carmen Morawetz, Angela R. Laird

## Abstract

Automated neuroimaging meta-analysis is widely used to generate hypotheses and interpret findings, reducing the time and effort required for manual literature searches and evidence curation. However, how closely these methods recover effortful expert-curated syntheses has not been established. We present NeuroMetaBench, an openly available benchmark linking expert curation records across stages of systematic review, from author-stated eligibility criteria and research-article inclusion decisions to individual analysis selections and meta-analysis-specific reference maps. The resource includes bibliographic records for 1,611 research-article inclusions across 17 quantitative review articles and candidate search pools for three. For nine review articles, it also provides analysis-level selections and coordinate data across 35 meta-analyses, documenting which reported results contributed to each meta-analysis and enabling reproducible construction of standardized reference maps. We demonstrate the benchmark’s utility by evaluating three established automated methods: Neurosynth, NeuroQuery, and NeuroVLM. Across the benchmark, automated maps generally showed limited to moderate spatial correspondence with expert-curated reference maps, with substantial variation across meta-analysis topics and characteristics. NeuroMetaBench provides an openly available, community-driven resource for evaluating established automated meta-analysis methods, and is well positioned to evaluate emerging LLM-based approaches to automated neuroimaging synthesis.

## Introduction

The neuroimaging literature continues to expand, with tens of thousands of studies published every year. Quantitative synthesis is essential both to integrate this growing literature and to address the limited statistical power of many individual studies (Poldrack et al., 2017). Researchers therefore routinely conduct coordinate-based meta-analyses (CBMA), systematically screening publications and pooling reported coordinates to identify consistent spatial patterns across studies of psychological constructs and phenotypes (Samartsidis et al., 2017; Wager et al., 2009). Nevertheless, producing a high-quality systematic meta-analysis remains time-consuming (Borah et al., 2017; Müller et al., 2018), limiting researchers’ ability to keep pace with an increasingly diverse and growing neuroimaging literature.

These challenges have motivated the development of automated methods that synthesize the neuroimaging literature at scale using natural language processing (NLP) techniques. Introduced in 2011, Neurosynth pioneered large-scale automated synthesis by combining text mining with coordinate extraction to map associations between psychological terms and brain activation (Yarkoni et al., 2011). Subsequent tools extended these capabilities through predictive modeling and learned representations of text and neuroimaging coordinates, enabling more flexible and fine-grained mapping from natural-language semantics to brain maps (Dockès et al., 2020; Ghayem et al., 2026; Hammonds et al., 2026; Meudec et al., 2024). Emerging large language model (LLM) workflows promise to extend this automation to reviews tailored to explicit research questions and eligibility criteria, with early biomedical applications reporting promising performance in article screening and structured evidence extraction (Sanghera et al., 2025; Zabaleta & Lin, 2026).

However, even as the use of automated syntheses continues to proliferate, systematic evaluation of their correspondence with expert neuroimaging meta-analyses remains limited. Existing evaluations have largely relied on comparisons with independent reference datasets, such as the Human Connectome Project (Yarkoni et al., 2011), and selected manually curated meta-analyses (Dockès et al., 2020). As a result, it is unclear how closely novel automated meta-analyses correspond to expert-curated equivalents. Outside neuroimaging, shared benchmarks offer precedents for evaluation: SYNERGY and SciLitBench provide expert-curated records of article screening decisions and, in the latter case, structured information extraction (De Bruin et al., 2023; Zabaleta & Lin, 2026). Benchmarks such as these enable researchers to evaluate automated literature review methods step-by-step across a wide range of review topics. Within neuroimaging, although there have been nascent efforts to systematically evaluate automated methods in structured metadata extraction (Sulaimanov et al., 2026) and reporting completeness (Pluimer et al., 2026), there does not yet exist a benchmark to evaluate automated meta-analysis end-to-end.

Moreover, these resources do not capture a key requirement of high-quality neuroimaging meta-analysis: selecting the specific analyses within eligible articles that address a target question. For instance, a meta-analysis of brain volume changes in schizophrenia must distinguish contrasts reporting lower volume in patients than controls (*controls > patients)* from those reporting the reverse effect (*patients > controls).* Such decisions directly impact the final inputs to the quantitative meta-analysis and depend on the populations, tasks, comparison conditions, and hypotheses targeted by a researcher. Commonly used meta-analytic coordinate formats also offer limited support for preserving unambiguous bibliographic links between reported analyses and publications, falling short of the standards required of a formal benchmark. As AI-based synthesis frameworks continue to evolve, it is becoming increasingly important to establish ground-truth benchmarks linking expert review criteria, research-article inclusion decisions, and analysis selections to quantitative reference maps, enabling evaluation of both the evidence selected and the synthesis it produces.

Here we present NeuroMetaBench, an openly available benchmark linking expert curation records across stages of neuroimaging meta-analysis. A primary goal is to establish a common, machine-readable structure linking review criteria and search information to research-article inclusion decisions, individual analysis selections, and quantitative reference maps. The initial release of the benchmark contains nine quantitative review articles reporting 35 meta-analyses comprising over 2,000 analyses from over 1,000 research-article records. In what follows, we describe NeuroMetaBench’s terminology and structure, as well as the tooling supporting community-led expansion into new topics. We then examine the nine review articles with analysis-level records, demonstrating that across meta-analysis topics there are substantial differences in research-article and analysis selection characteristics. Finally, we demonstrate the utility of our benchmark by evaluating three established automated frameworks—Neurosynth, NeuroQuery, and NeuroVLM— and explore how differences in performance across topics may relate to the selection characteristics of the expert references.

## Methods

### Dataset and Curation

The current collection assembles materials from published review articles in neuroimaging, including author-stated eligibility criteria and search information, bibliographic inclusion lists, candidate search pools, and author-supplied records of selected analyses and coordinates. It constitutes an initial convenience sample determined by the availability of recoverable expert records. The initial dataset was curated by the authors through iterative manual and semi-automated review of the recovered records.

### Terminology and record structure

NeuroMetaBench represents curation across three levels: the Protocol, Research-article inclusion, and Analysis Records (Table 1). Stable identifiers link records across these levels while bibliographic identifiers allow publications to be linked to public repositories.

**Table 1.** Coverage of the three curation tiers

| Tier | Content | Coverage |
|---|---|---|
| A Protocol | Review question, search information, eligibility criteria, record counts, method, and modality | 22 registered review articles<br>14 complete protocols |
| B Inclusion | Final Lists of included research articles linked by publication identifiers | 17 review articles<br>1,611 research-article inclusions<br>3 candidate pools |
| C Analysis | Analyses, coordinates, meta-analysis membership assignments, and reference-map products | 9 review articles<br>35 meta-analyses |

### Protocol

The registry defines the available review articles and assigns each a review-article identifier based on the PMID and/or DOI of the review article. It also records the authors’ inclusion and exclusion criteria and PubMed search criteria as stated in the original publication.

### Research-article inclusion

The included-research-article table links included research articles to review-article identifiers, using PMID as the primary research-article identifier and DOI as a backup or alternative. These links preserve review-specific inclusion decisions while allowing research articles shared across review articles to be identified.

Where authors provide lists of candidate research articles, a candidate-research-article table links these records to their respective review articles and records inclusion or exclusion decisions and associated reasons where available. Missing decisions or reasons are retained as missing rather than inferred. Note that review articles including lists of candidate research articles or search records are in the minority.

### Analysis records

An analysis is an individual statistical result from a contributing research article, represented by its reported coordinate set. In typical fMRI research articles, an analysis corresponds to a statistical *contrast*, although not every reported result is a contrast. An analysis may contribute to one or more **meta-analyses** reported in a review article.

Records use the Neuroimaging Meta-Analysis Data Standard (NiMADS), a machine-readable specification that supports interoperable representation of neuroimaging meta-analytic data (Kent et al., 2026). A NiMADS dataset groups analyses within study records, which correspond here to research articles, with each analysis representing an individual statistical result and its associated coordinates. Analysis records retain names, descriptions, and experimental conditions where available; coordinate records specify three-dimensional locations and stereotactic space. Research-article metadata can capture bibliographic information, sample sizes, and participant characteristics. NiMADS also supports links to statistical images and analysis-level annotations. This structure preserves distinctions among results reported within the same publication and supports their selection for meta-analytic estimation through NiMARE (Salo et al., 2023).

We refer to meta-analyses reported within the same review article as sibling meta-analyses; these may draw on overlapping or distinct sets of research articles and analyses.

### Meta-analysis membership annotations

To preserve complete provenance of analysis selection, NeuroMetaBench records analyses separately from their meta-analysis membership assignments, using an accompanying NiMADS annotation file for each review article. This distinction allows reported analyses to contribute to one or more meta-analyses, while representing the coordinates of each analysis only once. In addition, this format supports the encoding of all reported analyses, including those not relevant to any meta-analysis; however, the initial collection primarily contains analyses assigned to at least one meta-analysis.

### Reference maps

The dataset includes meta-analysis-specific reference maps generated from the corresponding expert-selected analyses. Each map is linked to its review article and the annotation defining its contributing analyses, preserving the connection between the curated evidence and the resulting quantitative synthesis. These maps provide reference outputs for evaluating automated methods (see Reference map generation). Reference-map products use multilevel kernel density analysis (MKDA), applying a common estimator across review articles that used different synthesis methods. The reference therefore represents synthesis of the expert-selected evidence under a standardized procedure, and not a precise numerical reproduction of the original published map. This separation allows the benchmark to preserve expert curation while making the map-generation procedure explicit and reproducible (Salo et al., 2023; Wager et al., 2007).

### Importing analysis records

NeuroMetaBench supports importing existing curation records using NiMARE utilities that support conversion from formats such as Sleuth into NiMADS. However, Sleuth records rely on potentially ambiguous author-based labels rather than structured bibliographic identifiers such as PMIDs, and do not explicitly group analyses by publication or link corresponding analyses across meta-analysis-specific files. The NeuroMetaBench repository therefore includes tools that combine fuzzy matching with human review: matching proposes candidate publications for each converted entry, and reviewers confirm or correct these associations before research-article identifiers are assigned. This process establishes publication provenance and supports consistent grouping of analyses within research articles. Corresponding analysis records are also checked across meta-analyses to preserve shared analyses and their meta-analysis membership assignments.

## Evaluation

### Reference map generation

Reference maps were generated for 35 meta-analyses from their expert-selected coordinates using NiMARE version 0.21.0. Meta-analyses were conducted using multilevel kernel density analysis (MKDA) (Wager et al., 2007), using a spherical kernel with a 10-mm radius and equal weighting of analyses. Each analysis contributed a binary indicator map, so overlapping spheres within an analysis counted only once at each voxel. Before analysis, we transformed 4,790 coordinates from Talairach to MNI space using the Lancaster transform (Lancaster et al., 2007). Maps were generated on the 2-mm MNI152 grid within an MNI152 brain mask. Each analysis was treated as an independent contribution; analyses from the same research article or overlapping participant samples were not combined or down-weighted.

Statistical inference used NiMARE’s approximate, histogram-based null distribution, with Benjamini–Hochberg false discovery rate (FDR) correction at q = 0.05 (Benjamini & Hochberg, 1995). Unthresholded z maps were used for spatial comparisons. FDR-corrected maps were used for thresholded displays and the reference eligibility criterion described below.

### Meta-analysis selection and eligibility

Four substance-use references—cannabis, nicotine, opioids and stimulants—were excluded because no voxels survived FDR correction, leaving 31 evaluable references. The spatial embedding in Figure 2 and identification analysis in Figure 4 additionally excluded the five ‘all’ meta-analyses that pool nearly all analyses from their review article (executive function, social cognition, problem solving, dementia and substance use), leaving 26 references. These broad meta-analyses were excluded to limit comparisons with near-complete pools of their own sibling meta-analyses. Two meta-analyses pooling the analyses included in sibling meta-analyses — emotion regulation reappraisal (containing every decrease and increase analysis) and cue reactivity reward (containing every drug- and natural-cue analysis) — were retained and marked separately.

**Figure 1.**
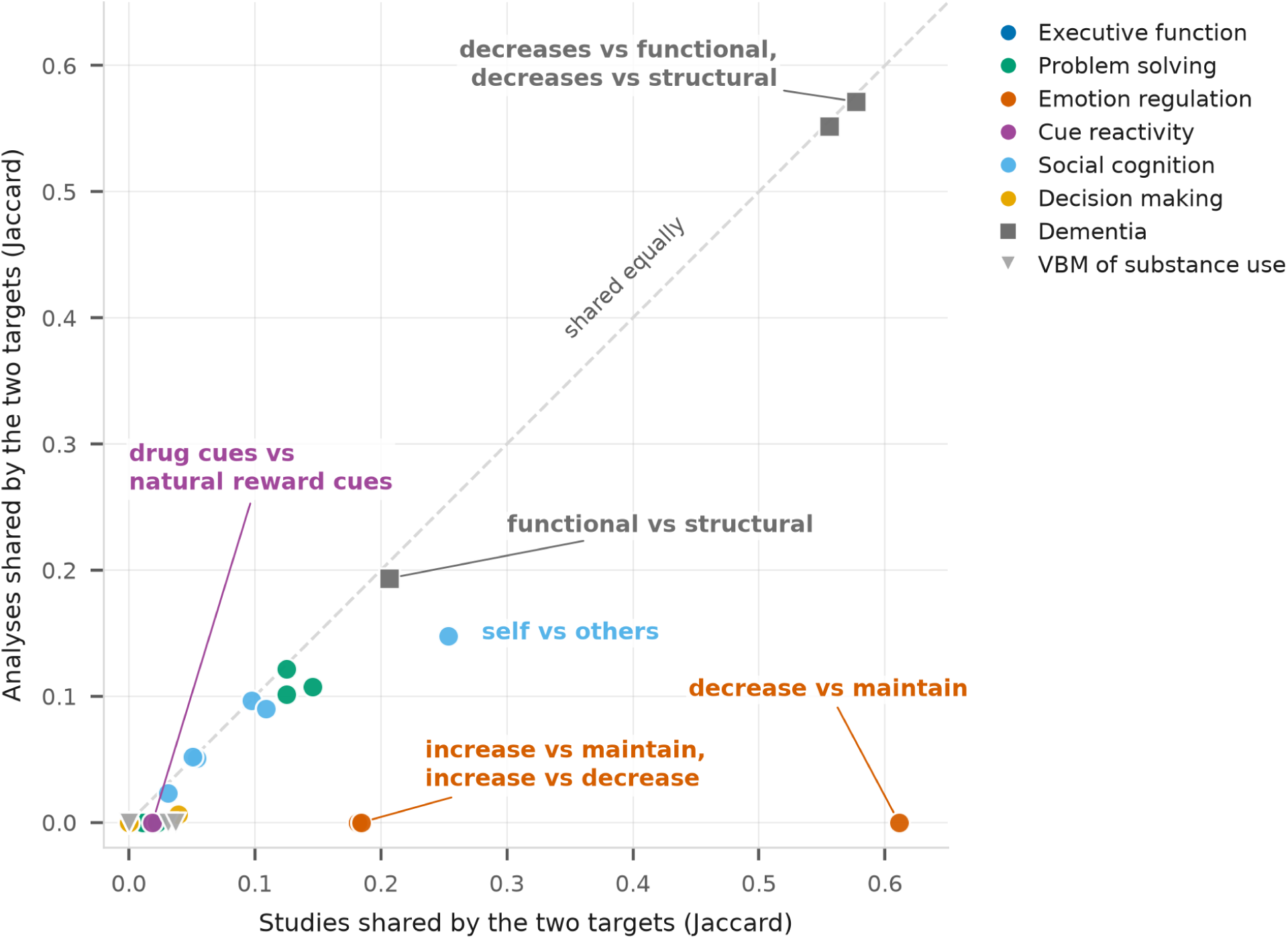
Research-article and analysis overlap between sibling meta-analyses. Each point represents a pair of meta-analyses within the same review article. The horizontal and vertical axes show the Jaccard indices for overlap in contributing research articles and analyses, respectively. Colour indicates review article. The dashed diagonal denotes equal overlap at both levels; points below the diagonal indicate lower analysis-level than research-article overlap. The five ‘all’ meta-analyses, containing ≥95% of their review article’s analyses or supersets of multiple narrower meta-analyses were excluded because their broad coverage would produce overlap by construction. The comparison included 35 pairs of selected meta-analyses.

**Figure 2.**
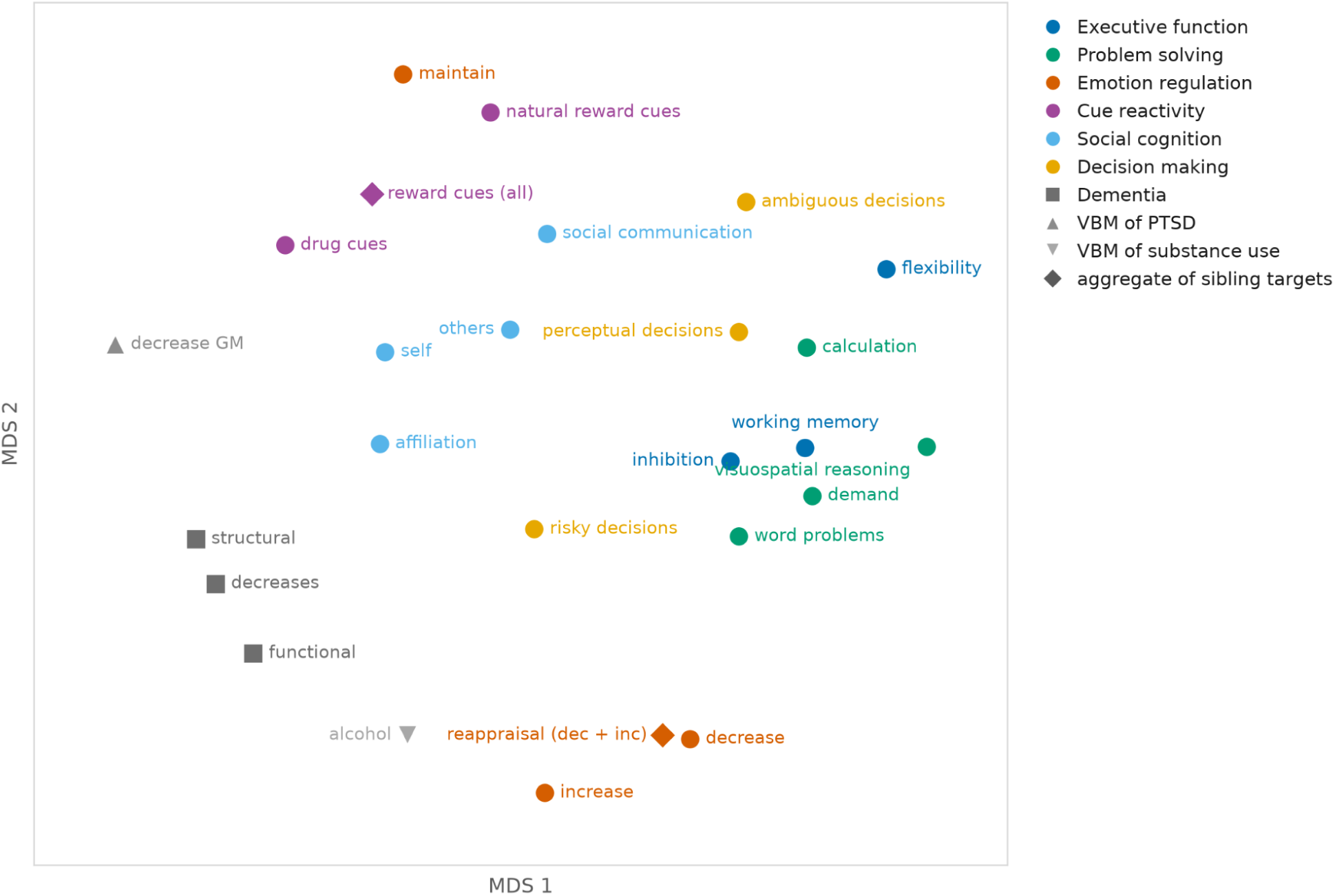
Spatial relationships among reference maps. Metric multidimensional scaling (MDS) of 26 reference maps, using correlation distance (1 − r) between unthresholded z maps within the brain mask. Nearby points represent more similar spatial patterns; axes are unitless, and only relative distances are meaningful. Colour indicates review article; squares denote dementia meta-analyses and triangles denote structural (VBM) meta-analyses. Meta-analyses containing ≥95% of their review article’s analyses were excluded: executive function all, social all, problem solving global, dementia all and substance use all drug classes. Diamonds denote retained meta-analyses that pool analyses included in sibling meta-analyses: emotion regulation reappraisal includes every decrease and increase analysis, and cue reactivity reward includes every drug- and natural-cue analysis. Maps correlated more strongly within review articles than between review articles (mean r = 0.51 versus 0.21). The MDS representation preserved between-map dissimilarities with a Shepard correlation of 0.84 and local neighbourhoods with a trustworthiness of 0.89 at k = 5.

**Figure 3.**
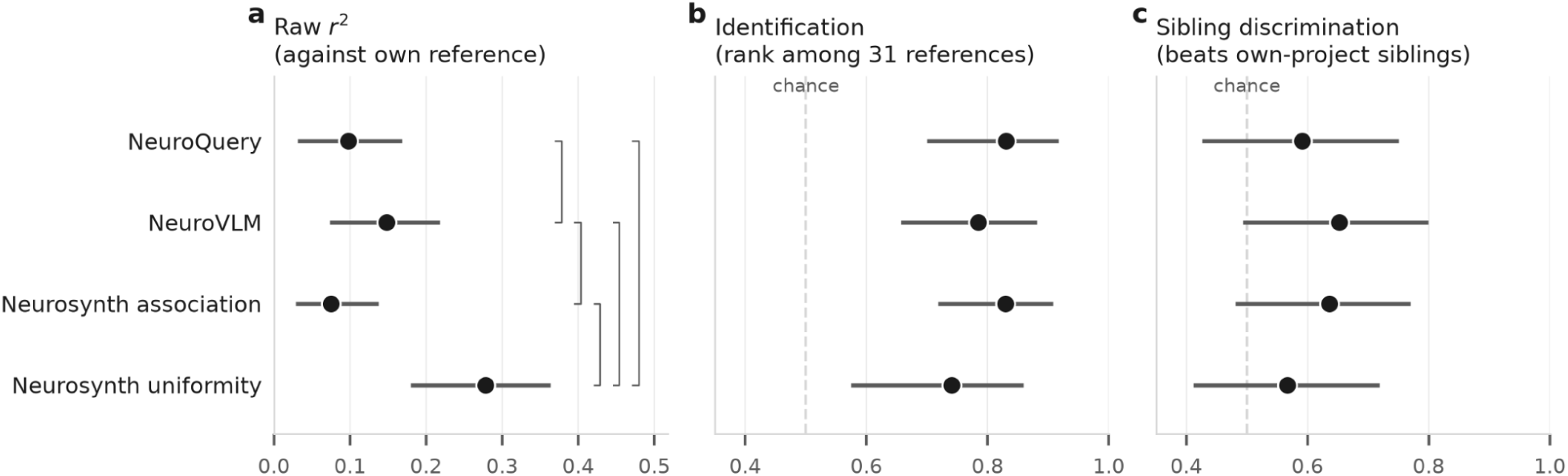
Performance of automated methods relative to expert-curated references. Values are means across 31 evaluable meta-analyses for Neurosynth association, NeuroQuery, NeuroVLM, and Neurosynth uniformity; panel c includes 30 meta-analyses with at least one sibling meta-analysis. Error bars indicate 95% confidence intervals estimated using a two-stage cluster bootstrap, resampling the nine review articles and then meta-analyses within each sampled review article (10,000 draws). (a) Squared voxelwise correlation with the intended reference. (b) Identification, defined as the proportion of the other 30 references with which an automated map correlated less strongly than with its intended reference. (c) Sibling discrimination, calculated using the same definition but restricted to alternative meta-analyses within the same review article. Dashed lines in panels b and c indicate chance performance (0.50). Brackets indicate statistically significant pairwise differences after Holm correction, based on exact sign-flip tests of mean differences calculated within each review article.

**Figure 4.**
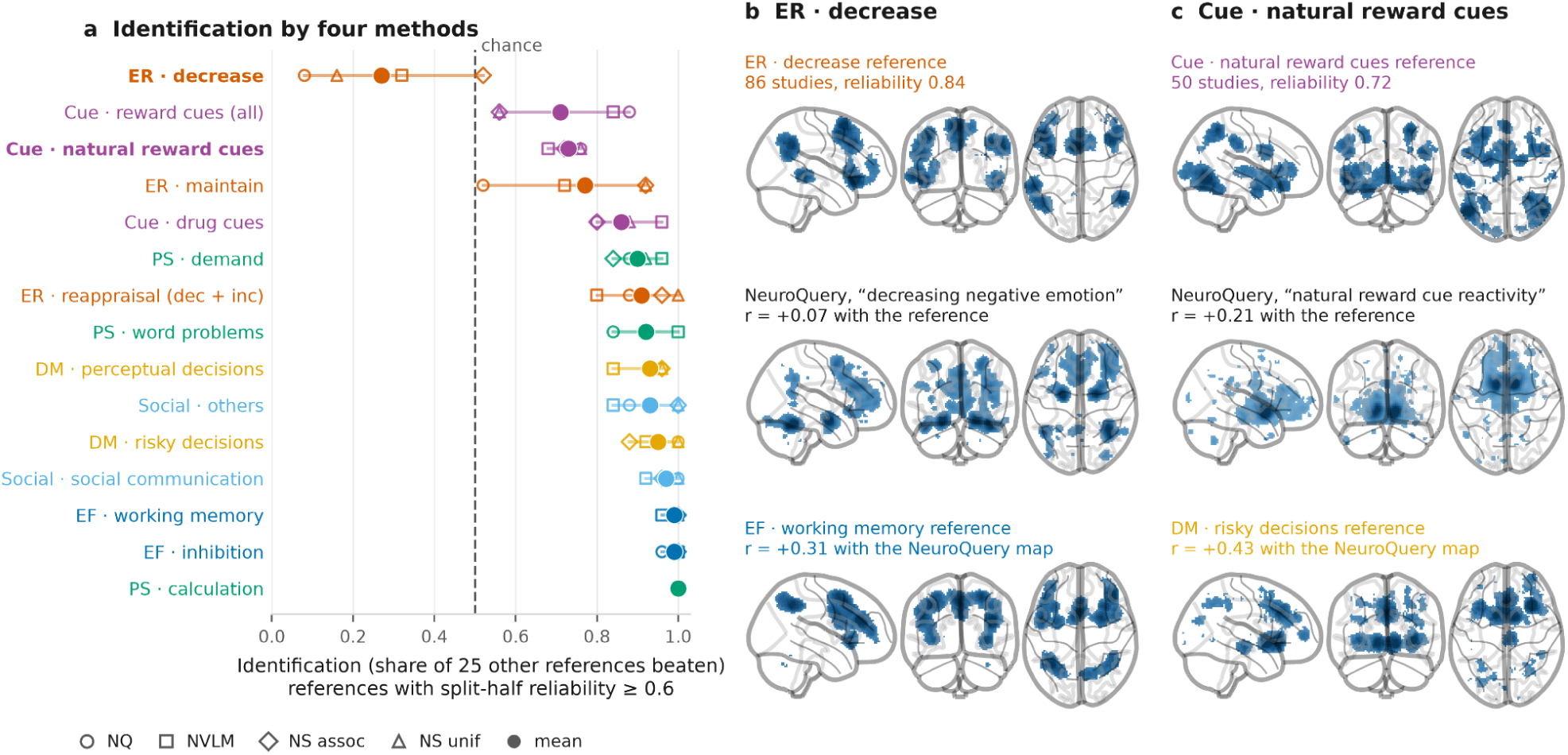
Automated methods can misidentify reliable reference maps. (a) Identification scores for NeuroQuery (NQ), NeuroVLM (NVLM), and Neurosynth association and uniformity maps. Scores indicate the proportion of alternative references with which an automated map correlates less strongly than with its intended reference (chance, 0.5). Only the 15 reference maps with split-half reliability ≥0.6 are shown. (b, c) Examples of confusion between meta-analyses, showing the intended reference, the NeuroQuery prediction and the alternative reference most strongly correlated with that prediction. (b) Emotion regulation decrease (86 research articles; split-half reliability, 0.84). The prediction for “decreasing negative emotion” correlates weakly with the intended reference (r = 0.07) and more strongly with executive function working memory (r = 0.31). (c) Cue reactivity natural cues (50 research articles; split-half reliability, 0.72). NeuroQuery’s prediction for “natural reward cue reactivity” correlates more strongly with the risky decision-making reference (r = 0.43) than with the intended reference (r = 0.21). The two meta-analyses share no research articles. Glass-brain displays show the top 5% of positive in-brain voxels in each map. All correlations are voxelwise Pearson correlations between unthresholded maps within the brain mask.

### Similarity structure of the reference maps

We calculated pairwise voxelwise Pearson correlations between the 26 unthresholded reference maps within the brain mask and converted them to correlation distances (1 − r). Mean correlations were summarized separately for pairs within and between review articles. The distance matrix was embedded in two dimensions using metric multidimensional scaling (MDS; scikit-learn version 1.9.1; (Pedregosa et al., 2012), implemented with SMACOF and 64 random initializations. A fixed random seed was used, and the solution with the lowest stress was retained.

### Reference map reliability

Reliability was estimated by repeatedly dividing each meta-analysis’s contributing research articles into two disjoint halves. All eligible analyses from a publication remained together, preventing the same publication from contributing to both halves. Where the number of research articles was odd, the additional research article was assigned to the second half. This procedure does not rule out participant overlap across separate publications.

An MKDA map was generated from each half using the reference-map settings, and the two unthresholded z maps were correlated within the brain mask. Each split-half correlation was adjusted using the Spearman–Brown formula, *2r/(1 + r)*, to estimate reliability at the full research-article count. We performed ten random splits per meta-analysis and summarized reliability by the mean corrected correlation, with the range across splits indicating sensitivity to the partition. References with mean reliability ≥0.6 were classified as reliable for Figure 4.

### Automated methods and query specification

We evaluated four map outputs from three methods: Neurosynth association and uniformity maps, NeuroQuery predictions, and NeuroVLM text-to-brain maps (Dockès et al., 2020; Hammonds et al., 2026; Yarkoni et al., 2011). Neurosynth maps were recomputed from the version 7 corpus of 14,371 research articles. Before fitting, all 464 research articles from the curated source records present in the corpus, including the 452 benchmark research articles, were removed, leaving 13,907 research articles.

For each Neurosynth term, research articles with an abstract tf-idf weight greater than 0.001—the default threshold—were contrasted with the remaining corpus using MKDA chi-squared with a prior of 0.5. This produced association maps representing reverse inference and uniformity maps representing forward inference. Infinite z values arising from p-value underflow were capped at the largest finite absolute z value in the map, preserving their sign. NeuroQuery maps were generated using neuroquery version 1.1.0 and its distributed pretrained model. NeuroVLM maps were generated using the text-to-brain pipeline with the Hugging Face models neurovlm/NeuroAutoEncoder (revision b7c288b) and neurovlm/ProjectionHeads (revision bcb7e8c). Model files were downloaded on 9 September 2026.

One query was written for each meta-analysis to express the construct defined by the source review article. NeuroQuery and NeuroVLM received free-text queries; Neurosynth terms were selected from its fixed vocabulary using the same queries and were fixed before any Neurosynth maps were computed. The 35 meta-analyses mapped to 29 distinct Neurosynth terms: 31 meta-analyses used a term appearing directly in the query, one used the vocabulary form of a query word, and three used a broader term because no meta-analysis-specific term was available. Consequently, some sibling meta-analyses shared a Neurosynth term. Complete queries, terms, model specifications and inference settings are provided in the Supplementary Materials.

### Spatial correspondence and identification

We evaluated how closely each automated method reproduced the spatial pattern of the expert-curated reference for the same meta-analysis, and whether its output matched that reference more strongly than alternative references. Automated maps were resampled to the reference maps’ 2-mm MNI152 grid using continuous interpolation in Nilearn (Nilearn contributors et al., 2026) and all comparisons used the same 228,483-voxel brain mask. Correspondence with the intended reference was measured as the squared voxelwise Pearson correlation (r²) between unthresholded maps.

To assess specificity to the intended meta-analysis, we calculated two additional metrics using signed Pearson correlations. The identification score was the proportion of alternative references with which an automated map correlated less strongly than with its intended reference. A score of 0.5 corresponds to chance, and a score of 1 indicates that the intended reference outranks every alternative. Sibling discrimination applied the same calculation to alternative meta-analyses within the same review article, testing whether an automated map distinguished closely related synthesis questions. This metric was undefined for the VBM PTSD review article, which contributed one meta-analysis.

### Aggregation and statistical inference

We summarized each output’s correspondence, identification, and sibling discrimination scores as unweighted means across eligible meta-analyses. To account for dependence within review articles, 95% confidence intervals were estimated using a two-stage cluster bootstrap: review articles were sampled with replacement, followed by meta-analyses within each sampled review article. We used 10,000 resamples and percentile intervals. Pairwise differences between automated outputs were tested using exact sign-flip permutation tests on mean differences calculated within each review article, with Holm correction across the six pairs for each metric (Holm, 1979). Agreement among the four automated outputs in their rankings of identification scores was summarized using Kendall’s W without tie correction, calculated across the 15 meta-analyses shown in Figure 4a.

## Data availability

NeuroMetaBench data and code are openly available at https://github.com/neurostuff/neurometabench, release *v*1.0, under the MIT license. Counts reported reflect all meta-analyses in the v1 inventory, released on November 20 2026. The benchmark is archived on Zenodo (de la Vega et al., 2026).

## Code availability

Code for the analyses in this publication is available at https://github.com/adelavega/nmb-paper under the MIT license.

## Results

### Composition of the analysis-level collection

The analysis-level collection comprised nine review articles spanning multiple research topics and imaging modalities (Table 3). Together, these records spanned 1,093 research-article records containing 2,300 analyses, 23,723 coordinates, and 35 meta-analyses. Review articles contributed between one meta-analysis, for voxel-based morphometry (VBM) of post-traumatic stress disorder (PTSD), and six meta-analyses, for VBM of substance use.

**Table 2.** Terminology and definitions of core NeuroMetaBench entities

| **Entity** | **Definition** |
|---|---|
| Quantitative review article | A publication supplying review criteria, curation records and one or more meta-analyses to the benchmark; shortened to review article after first use. |
| Research article | A primary publication screened for inclusion, potentially reporting multiple analyses. |
| Analysis | An individual statistical result from a contributing research article, represented by its reported coordinate set. |
| Meta-analysis | An individual synthesis combining eligible analyses to address a particular question and produce a spatial map. |
| Contrast | A specific statistical comparison. |

**Table 3.** Composition of the analysis-level collection

| Review article | Modality | Research-article records | Analyses | Coordinates | Meta-analyses | Analyses per research article |
|---|---|---|---|---|---|---|
| Cue reactivity | fMRI | 191 | 275 | 3,270 | 3 | 1.44 |
| Decision making | fMRI | 136 | 225 | 1,955 | 3 | 1.65 |
| Dementia | fMRI, Structural | 29 | 31 | 1,410 | 4 | 1.07 |
| Emotion regulation | fMRI | 90 | 248 | 2,829 | 4 | 2.76 |
| Executive function | fMRI | 178 | 271 | 3,076 | 4 | 1.52 |
| Problem solving | fMRI | 129 | 282 | 3,166 | 5 | 2.19 |
| Social cognition | fMRI | 238 | 856 | 7,185 | 5 | 3.60 |
| VBM of PTSD | Structural | 22 | 25 | 159 | 1 | 1.14 |
| VBM of substance use | Structural | 80 | 87 | 673 | 6 | 1.09 |
| **Total** |  | **1,093** | **2,300** | **23,723** | **35** | **2.10** |

Sources: Cue reactivity (Hill-Bowen et al., 2021); Decision making (Poudel et al., 2020); Dementia (Kamalian et al., 2022); Emotion regulation (Morawetz et al., 2022); Executive function (Niendam et al., 2012); Problem solving (Bartley et al., 2018); Social cognition (Pintos Lobo et al., 2023); VBM of PTSD (Pankey et al., 2022); VBM of substance use (Hill-Bowen et al., 2022).

Review articles varied in how their constituent research articles contributed to specific meta-analyses. Across the collection, 49% of research articles contributed more than one annotated analysis (mean, 2.10 analyses per research article; range, 1–13). However, many meta-analyses within the same review article drew on largely distinct literatures, whereas others shared contributing publications but selected different analyses from them. To characterize these patterns, we calculated pairwise Jaccard overlap between meta-analyses within each review article, separately at the research-article and analysis levels (Figure 1).

The emotion regulation review article illustrated how shared publications can support distinct meta-analysis-specific selections, with 70% of research articles contributing to more than one meta-analysis. Its decrease, increase, and maintain meta-analyses had a mean research-article overlap of 0.33 but shared no individual analyses. The decrease and maintain meta-analyses, for example, shared 55 of the 59 research articles contributing to maintain while selecting entirely different contrasts. In contrast, review articles on executive function, VBM of substance use, decision making and cue reactivity showed little overlap between meta-analyses at either level (mean research-article Jaccard index ≤0.02). For instance, in the executive function review article, the working-memory, inhibition, and flexibility meta-analyses had maximum pairwise research-article Jaccard indices of 0.02. Similarly, in VBM of substance use, each meta-analysis represented a different drug class and drew largely on different research articles. In cue reactivity, the drug-cue and natural-cue meta-analyses shared only 3 of their 160 research articles. Social cognition and problem solving showed intermediate research-article overlap (mean Jaccard indices, 0.10 and 0.07, respectively), whereas dementia meta-analyses overlapped substantially at both the research-article and analysis levels (0.45 and 0.44, respectively).

### Spatial relationships and reliability of reference maps

The reference meta-analytic maps exhibited diverse spatial patterns (Figure 2). Mean voxelwise correlation was higher within review articles than between review articles (0.51 versus 0.21), indicating greater spatial similarity among related meta-analyses. In the multidimensional scaling representation, executive function, problem solving and perceptual decision making occupied neighbouring regions, whereas dementia maps clustered apart from the task-based maps. Emotion regulation ‘maintain’ lay closer to cue reactivity meta-analyses than to other emotion regulation meta-analyses, while the reappraisal aggregate lay beside the decrease meta-analysis whose analyses it includes.

We estimated the split-half reliability of reference spatial maps across ten random splits (Figure S1) and observed substantial variability across review articles. Most meta-analyses showed moderate to high reliability, with 21 of the 31 yielding a mean Spearman–Brown-corrected correlation ≥0.5. Examples of high reliability included executive function working memory and inhibition (0.72–0.85) and emotion regulation decrease (0.84). Risky decision making also showed moderate reliability (0.65; 65 research articles). The least reliable meta-analyses were VBM of PTSD (0.25; 22 research articles), VBM of alcohol use (0.27; 21 research articles), dementia functional findings (0.33; 17 research articles), and ambiguous decision making (0.38; 41 research articles). Across these four meta-analyses, the halves contained 8–21 research articles.

However, meta-analyses from the two structural VBM review articles showed low reliability across a range of research-article counts, including the substance-use all-drug-classes meta-analysis (0.30) despite its 80 research articles.

### Correspondence between automated methods and expert-curated references

Next, we estimated how well three automated methods—Neurosynth, NeuroQuery, and NeuroVLM—could recover the reference meta-analytic maps. Spatial correspondence varied substantially across outputs (Figure 3), with mean squared voxelwise correlations (r²) of 0.075 for Neurosynth association, 0.098 for NeuroQuery, 0.148 for NeuroVLM, and 0.279 for Neurosynth uniformity. The higher correspondence of Neurosynth uniformity maps is consistent with their closer alignment to conventional coordinate-based meta-analysis: uniformity summarizes where activation is reported for a term rather than identifying term-specific associations or predictive regions. However, spatial correspondence can also reflect activation in regions shared across tasks, without capturing what distinguishes the intended meta-analysis.

To assess specificity to the intended meta-analysis, we measured how often each automated map correlated more strongly with its intended reference than with other maps. The four outputs performed similarly on this “identification” measure, with mean scores of 0.74–0.83 against a chance level of 0.50 and no significant pairwise differences. Neurosynth uniformity had the lowest mean identification score despite its advantage in spatial correlation, suggesting that greater overall correspondence did not translate into better discrimination between meta-analyses.

Distinguishing closely related meta-analyses proved more difficult for all methods. When identification was restricted to sibling meta-analyses within the same review article, mean discrimination scores fell to 0.57–0.65, and every method’s confidence interval included chance. Thus, although automated maps generally distinguished their intended references from the broader collection, they showed limited ability to resolve finer distinctions among questions within a research topic. Restricting the comparison to 18 meta-analyses with split-half reliability ≥0.6 yielded mean discrimination scores of 0.61–0.71. All confidence intervals included chance, although the lower bound for Neurosynth uniformity was exactly 0.50. These results indicate that difficulty distinguishing related meta-analyses persisted among reliable references.

### Automated performance varied across review articles

Aggregate scores concealed substantial variation across meta-analyses. Among references with high split-half reliability (≥0.6), mean identification scores across the four outputs ranged from 0.27 to 1.00 (Figure 4a). The approaches broadly agreed on which meta-analyses were difficult to identify (Kendall’s W = 0.68), indicating that identification failures were shared across methods and were not confined to references with unstable spatial patterns.

Emotion regulation “decrease” provided a striking example (Figure 4b). Despite high reference reliability (0.84), NeuroQuery’s prediction for “decreasing negative emotion” correlated only weakly with the intended reference (r = 0.07). It instead correlated most strongly with the executive function working-memory reference (r = 0.31). NeuroQuery’s prediction also highlighted regions, such as the amygdala and middle temporal gyrus, that were not present in the expert reference map. This failure occurred in a review article where meta-analyses drew heavily on a shared publication set but selected distinct analyses, illustrating the importance of preserving contrast-level distinctions.

Automated methods also confused meta-analyses drawn from distinct literatures. The natural-cue reactivity reference was reliable (split-half reliability, 0.72; 50 research articles), yet all four automated outputs correlated most strongly with the risky decision-making reference rather than their intended reference (Figure 4c). NeuroQuery’s prediction for “natural reward cue reactivity” correlated weakly with the natural-cue reference (r = 0.21) and more strongly with risky decision making (r = 0.43), yielding an identification score of 0.76. This mismatch occurred despite no overlap among the 115 research articles contributing to the two meta-analyses and only modest similarity between their reference maps (r = 0.25). NeuroQuery’s prediction emphasized striatal regions shared with the risky decision-making reference, whereas the natural-cue reference predominantly involved visual and attentional cortex. This pattern suggests that the automated maps captured a broad reward-related pattern more readily than the spatial pattern associated with the specific cue-reactivity question.

## Discussion

NeuroMetaBench is an open resource for benchmarking automated neuroimaging meta-analysis tools, linking expert review criteria and curation decisions to quantitative reference maps. Precise bibliographic records establish the provenance of selected evidence, allowing it to be traced across review articles. A common, machine-readable representation supports reproducibility by preserving the records needed to reconstruct each synthesis. This framework enables the evaluation of automated neuroimaging methods both at intermediate decisions and the final output syntheses. By bringing together meta-analyses across diverse neuroimaging topics, NeuroMetaBench also enables researchers to characterize how syntheses are constructed and identify opportunities to make future evidence synthesis more precise, reproducible, and scalable.

A distinctive contribution of NeuroMetaBench is its analysis-level record of which reported results contribute to each meta-analysis. These records revealed substantial variation in how review articles assembled their evidence. For instance, executive function, decision making, and VBM of substance use drew on largely separate literatures for each meta-analysis, whereas emotion regulation divided a shared literature by contrast. Review articles also differed in whether meta-analysis membership assignments were exclusive. In social cognition, individual analyses could contribute to multiple meta-analyses, reflecting the hypothesis that different tasks recruit overlapping cognitive processes (Pintos Lobo et al., 2023). We also observed that review articles differed in the specificity of their analysis-level inclusion rules: some admitted only particular contrasts, whereas others primarily excluded results based on disqualifying criteria (e.g. region-of-interest or connectivity analyses), allowing a wider range of contrasts to qualify. The wide range of patterns across these review articles highlights the importance of NeuroMetaBench’s detailed provenance record, as review-specific selection rules can yield entirely different evidence from the same publications.

These differences in evidence selection raise the question of whether existing automated methods can recover the specific syntheses generated by experts. Our evaluation of Neurosynth, NeuroQuery, and NeuroVLM suggests only a partial recovery. Spatial correspondence was broadly limited across all methods, with Neurosynth uniformity maps showing the highest correspondence, consistent with their closer alignment to conventional coordinate-based meta-analysis. This advantage did not extend to identification, with all four outputs performing similarly when asked to distinguish the intended reference from alternatives.

Performance declined further when alternatives were restricted to sibling meta-analyses, indicating that recognizing a broad topic was easier than recovering distinctions between related synthesis questions. Poor sibling identification also occurred for highly reliable reference maps, suggesting that instability in the expert maps alone cannot explain the observed limitations.

Performance varied substantially across review articles, offering clues to where automated synthesis falls short. Emotion regulation was a particularly striking case: its meta-analyses drew heavily on the same publications while selecting distinct contrasts, yet even highly reliable references were poorly identified across methods. This example highlights a potential limitation shared by the evaluated approaches: they associate article-level text with pooled reported coordinates without explicitly distinguishing individual analyses or selecting those relevant to a target question, as an expert would (Dockès et al., 2020; Hammonds et al., 2026; Yarkoni et al., 2011). This loss of contrast-specific information may limit how closely these methods can reproduce expert syntheses, even when they identify the relevant literature.

This constraint may matter less where meta-analyses represent broad distinctions supported by largely separate literatures. For instance, all evaluated methods identified executive function references relatively well. These meta-analyses drew on largely distinct sets of research articles and represented broad, well-established cognitive constructs, such as inhibition and working memory. Separate literatures did not guarantee successful discrimination, however. All four automated outputs for natural-cue reactivity most closely resembled the risky decision-making reference, despite the two meta-analyses sharing no research articles. The references themselves were only modestly correlated, and NeuroQuery’s prediction emphasized striatal regions shared with risky decision making rather than the predominantly visual and attentional pattern of the natural-cue reference. This suggests that broad reward-related associations may overshadow distinctions between specific research questions. Whereas emotion regulation highlights a potential limitation in selecting different analyses from shared publications, natural-cue reactivity illustrates a complementary difficulty in distinguishing questions supported by separate literatures. Neither comparison alone establishes whether the mismatch arises from query interpretation, evidence selection or map estimation.

Together, these examples suggest that existing automated methods can capture some broad distinctions in cognitive neuroscience but struggle with others, even when meta-analyses draw on different literatures. Executive function meta-analyses may be easier to identify because of distinct neural patterns, more consistent terminology across publications, or both. Future research and uses of NeuroMetaBench should focus on better understanding which factors predict whether a topic or construct can be captured using existing automated methods, and how to improve the methods themselves to better separate constructs that experts define as separable.

### Limitations and future directions

NeuroMetaBench links review criteria, research-article inclusion decisions, and analysis selections to quantitative reference maps, supporting evaluation of both selected evidence and the resulting synthesis. The present evaluation was limited to final maps because the evaluated methods did not expose a complete sequence of eligibility decisions and analysis-to-meta-analysis assignments. Consequently, the observed discrepancies cannot be attributed separately to research-article retrieval, query interpretation, contrast selection, or estimation. As automated workflows evolve and expose these intermediate decisions, NeuroMetaBench can support stage-by-stage evaluation to identify where errors arise and how they affect the final synthesis. This is particularly relevant to emerging LLM workflows for article screening (Sanghera et al., 2025) and multi-stage review tasks, including full-text assessment and structured evidence extraction (Zabaleta & Lin, 2026). The resource complements screening collections such as SYNERGY and SciLitBench (De Bruin et al., 2023; Zabaleta & Lin, 2026) as well as benchmarks of evidence interpretation (DeYoung et al., 2020), extraction (Nye et al., 2018; Sulaimanov et al., 2026) and narrative synthesis (DeYoung et al., 2021), by connecting neuroimaging-specific analysis selections to quantitative reference maps.

The current analysis-level collection is a convenience sample of nine review articles, determined by the availability of recoverable expert records and weighted toward task-based fMRI. As such, each record reflects the decisions of a particular researcher or team at a particular time, whose accuracy, consistency, and generalizability remain incompletely characterized. The original reviews were bounded by their search dates, sources, and eligibility criteria, which may have limited coverage of the relevant literature. In addition, records primarily cover analyses assigned to at least one meta-analysis, leaving results excluded from every meta-analysis largely undocumented. These limitations make it difficult to determine from the benchmark alone why some review articles yield more reproducible maps or are better recovered by automated methods. Resolving these questions will require broader efforts across cognitive neuroscience, including independent curation, new experiments, and more consistent terminology for describing tasks and cognitive constructs using community-driven ontologies such as Cognitive Atlas (Poldrack et al., 2011).

In addition, the current reference maps rely on coordinate-based meta-analysis, which retains only reported peak locations and discards much of the information available in full statistical images. Image-based meta-analysis (IBMA) preserves this information and can provide a more faithful synthesis of the contributing research articles (Salimi-Khorshidi et al., 2009). Infrastructure for sharing statistical maps is available through NeuroVault (Gorgolewski et al., 2015) and Neurosynth Compose supports image-based synthesis (Kent et al., 2026). NeuroMetaBench can accommodate image-based records in principle by linking to external repositories such as NeuroVault, enabling richer comparisons and reducing dependence on selectively reported coordinates. The present evaluation, however, is limited to coordinate-based references, and its findings should be interpreted within the constraints of that representation.

Taken together, NeuroMetaBench should be interpreted as a collection of context-dependent expert references rather than definitive ground truth about the underlying neurobiology. As an open, expandable resource, however, it offers researchers an opportunity to pool their meta-analytic contributions and build cumulative knowledge about how evidence is selected, how these decisions shape syntheses, and how automated methods can better recover the distinctions that matter for specific research questions. Broader community contributions can extend coverage across topics, modalities, and curation practices, while independent annotations, explicit exclusion records, author corrections, and updated searches can clarify uncertainty in those decisions. As an open resource of expert-curated decisions, NeuroMetaBench provides a common basis for evaluating automated synthesis as advances in natural language processing accelerate the proliferation of AI-generated evidence syntheses.

