## Supplemental Information for "NeuroMetaBench: Expert-curated benchmark for evaluating automated neuroimaging meta-analysis"

### NeuroMetaBench: Supplementary Information

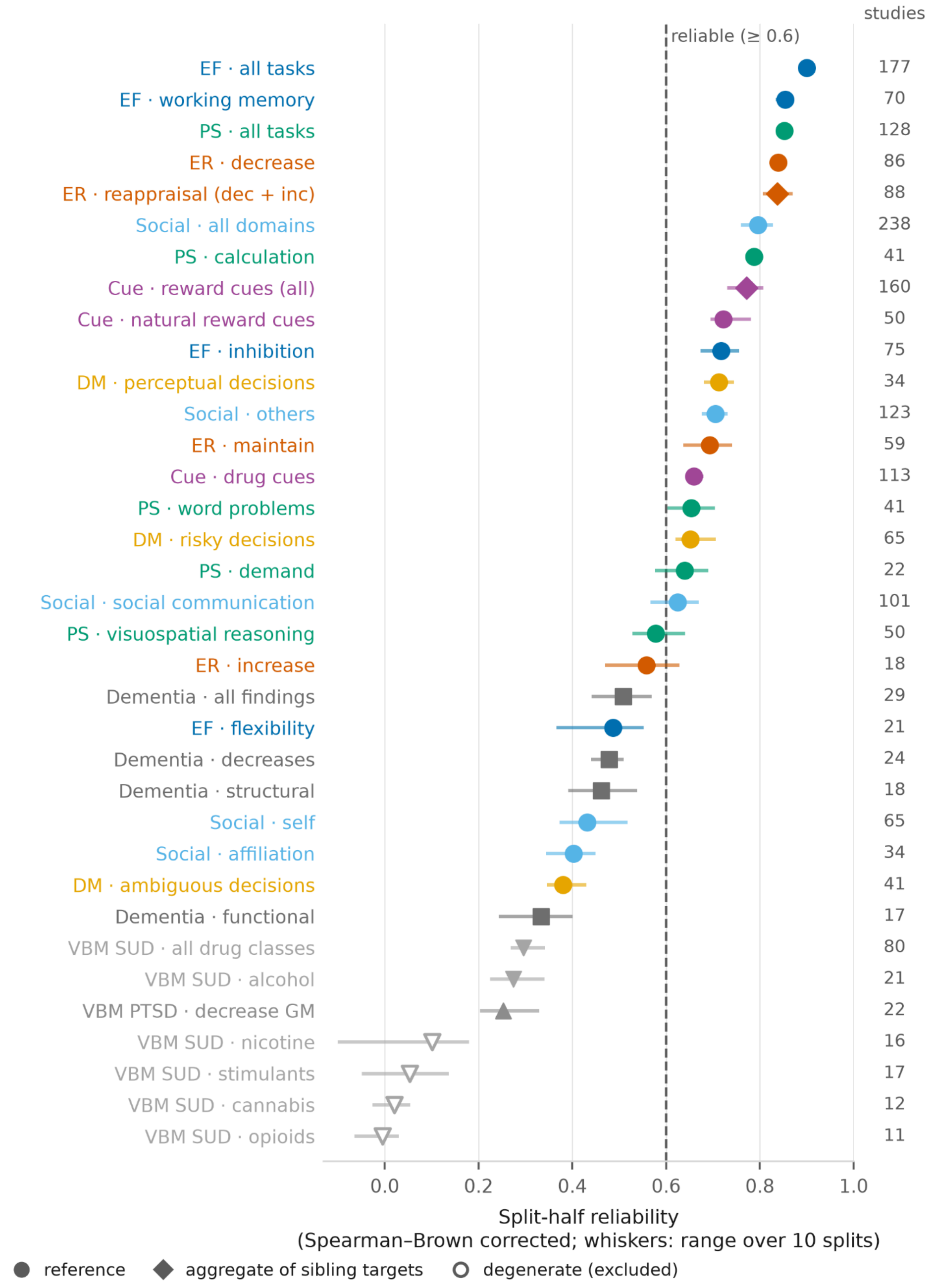


**Figure S1. Split-half reliability of the benchmark references.** Each row is one of the 35 benchmark meta-analyses, ordered by reliability: the Spearman–Brown corrected correlation between maps estimated from two disjoint halves of the contributing research articles, averaged over ten random splits (dot), with the range across splits (whiskers). The column at right gives the number of contributing research articles. Colour indicates review article; diamonds mark aggregates of sibling meta-analyses; hollow symbols mark the four degenerate references excluded from evaluation. The dashed line marks the 0.6 threshold used to define reliable references in Figure 4. Abbreviations: EF, executive function; PS, problem solving; ER, emotion regulation; DM, decision making; SUD, substance use disorder; GM, gray matter.

Table S1. Queries and Neurosynth terms for the 35 benchmark meta-analyses.

| **Review article** | **Meta-analysis** | **Research articles** | **Analyses** | **Reliability** | **Query (NeuroQuery, NeuroVLM)** | **Neurosynth term** | **Term source** |
|---|---|---|---|---|---|---|---|
| Cue reactivity | reward cues (all)a | 160 | 221 | 0.77 | cue reactivity | cue | direct |
| | drug cues | 113 | 137 | 0.66 | drug cue reactivity | drug | direct |
| | natural reward cues | 50 | 84 | 0.72 | natural reward cue reactivity | reward | direct |
| Decision making | ambiguous decisions | 41 | 58 | 0.38 | decision making under ambiguity | ambiguous | vocabulary form |
| | risky decisions | 65 | 103 | 0.65 | risky decision making | risky | direct |
| | perceptual decisions | 34 | 65 | 0.71 | perceptual decision making | perceptual | direct |
| Dementia | all findingsb | 29 | 31 | 0.51 | dementia | dementia | direct |
| | decreases | 24 | 26 | 0.48 | decreased brain activity in dementia | dementia | direct |
| | functional | 17 | 18 | 0.33 | functional activation in dementia | dementia | direct |
| | structural | 18 | 19 | 0.46 | gray matter atrophy in dementia | atrophy | direct |
| Emotion regulation | reappraisal (dec + inc)a | 88 | 154 | 0.84 | cognitive reappraisal | reappraisal | direct |
| | decrease | 86 | 128 | 0.84 | decreasing negative emotion | emotion | direct |
| | increase | 18 | 24 | 0.56 | increasing emotional | emotional | direct |

| | | | | | | | |
|---|---|---|---|---|---|---|---|
| | | | | | response | | |
| | maintain | 59 | 94 | 0.69 | maintaining emotional response | emotional | direct |
| Executive function | all tasksb | 177 | 259 | 0.90 | executive function | executive function | direct |
| | working memory | 70 | 124 | 0.85 | working memory | working memory | direct |
| | inhibition | 75 | 96 | 0.72 | response inhibition | response inhibition | direct |
| | flexibility | 21 | 23 | 0.49 | cognitive flexibility task switching | flexibility | direct |
| Problem solving | all tasksb | 128 | 281 | 0.85 | problem solving reasoning | reasoning | direct |
| | calculation | 41 | 100 | 0.79 | mental arithmetic calculation | calculation | direct |
| | word problems | 41 | 93 | 0.65 | verbal problem solving | verbal | direct |
| | visuospatial reasoning | 50 | 88 | 0.58 | visuospatial reasoning | visuospatial | direct |
| | demand | 22 | 41 | 0.64 | logical reasoning | reasoning | direct |
| Social cognition | all domainsb | 238 | 856 | 0.80 | social cognition | social cognition | direct |
| | affiliation | 34 | 106 | 0.40 | attachment and affiliation | social | broader |
| | self | 65 | 206 | 0.43 | self referential processing | self referential | direct |
| | others | 123 | 407 | 0.70 | person perception of others | person | direct |

| | | | | | | | |
|---|---|---|---|---|---|---|---|
| | social communication | 101 | 381 | 0.62 | social communication | communication | direct |
| VBM of PTSD | decrease GM | 22 | 25 | 0.25 | posttraumatic stress disorder gray matter | posttraumatic | direct |
| VBM of substance use | all drug classesb | 80 | 87 | 0.30 | substance use disorder gray matter | substance | direct |
| | alcohol | 21 | 22 | 0.27 | alcohol dependence gray matter | alcohol | direct |
| | nicotinec | 16 | 17 | 0.10 | nicotine smoking gray matter | smoking | direct |
| | opioidsc | 11 | 12 | -0.01 | opioid dependence gray matter | dependence | direct |
| | stimulantsc | 17 | 18 | 0.05 | stimulant use gray matter | substance | broader |
| | cannabisc | 12 | 12 | 0.02 | cannabis use gray matter | substance | broader |

Research articles and analyses: those contributing to each meta-analysis. Reliability: mean Spearman–Brown-corrected split-half correlation over ten random splits of the contributing research articles (Figure S1).

Query: the single free-text query written for each meta-analysis and given to NeuroQuery and NeuroVLM. Neurosynth term: selected from the version 7 vocabulary using the same query, fixed before any Neurosynth map was computed; sibling meta-analyses may share a term. Term source: direct, the term appears in the query; vocabulary form, the vocabulary's form of a query word; broader, no meta-analysis-specific term was available.

A, Pools the analyses of sibling meta-analyses (reappraisal: every decrease and increase analysis; reward cues: every drug- and natural-cue analysis). B, Contains ≥95% of its review article's analyses; excluded from Figures 2 and 4. cNo voxels survived FDR correction; excluded from evaluation.
